# Identification of a SARS-CoV-2 Spike RNA-Cleaving DNAzyme and Optimization of Its AS1411 Chimera

**DOI:** 10.64898/2026.09.01.748480

**Authors:** Zehra Banu Ünlü, Hüseyin Saygın Portakal, Osman Doluca

## Abstract

Sequence-specific cleavage of viral RNA by deoxyribozymes offers a direct approach for suppressing viral gene expression, although limited cellular accessibility remains a major barrier to their functional application. In this study, a panel of eight deoxyribozymes representing different catalytic motifs and target regions was designed against SARS-CoV-2 Spike RNA and comparatively evaluated through time-dependent in vitro cleavage assays. Substantial target-site-dependent differences were observed even among candidates sharing the same catalytic core, demonstrating that catalytic performance was strongly influenced by the selected RNA target region. Among the tested sequences, 8-17-769 exhibited the fastest and most reproducible cleavage profile and the highest observed rate constants, and was therefore selected as the catalytic module for subsequent chimera development. To evaluate its activity in a cellular context, 8-17-769 was combined with the nucleolin-binding aptamer AS1411 in two opposite linear orientations. In an A549-based Spike expression model, both AS1411-containing chimeras were associated with reduced S gene expression, with 8-17-769–AS1411 producing the strongest response, corresponding to an approximately tenfold reduction relative to the reference group and reaching statistical significance (p < 0.05). The functional difference between the two orientations was further examined structurally. Circular dichroism spectroscopy showed that 8-17-769–AS1411 more closely preserved the spectral characteristics of the individual components, while computational analyses indicated a dynamic organization more similar to the free deoxyribozyme–RNA complex and a nucleolin-compatible docking configuration. Collectively, these findings identify 8-17-769 as a lead Spike RNA-cleaving deoxyribozyme and demonstrate that the linear organization of aptamer and catalytic modules can substantially influence the structural and functional properties of aptamer–deoxyribozyme chimeras.

## INTRODUCTION

Coronavirus disease 2019 (COVID-19), caused by severe acute respiratory syndrome coronavirus 2 (SARS-CoV-2), rapidly developed into a worldwide pandemic following its emergence in late 2019 and created an unprecedented demand for effective antiviral strategies (Dash et al., 2021; Soares et al., 2025). SARS-CoV-2 is an enveloped betacoronavirus carrying a positive-sense single-stranded RNA genome of approximately 30 kb, in which viral replication and propagation depend on the coordinated expression of multiple viral genes (Brant et al., 2021; Chen et al., 2020). The RNA-based nature of the viral genome therefore provides an opportunity for sequence-specific therapeutic approaches designed to directly interfere with viral transcripts. Although the rapid development of vaccines represented a major milestone in controlling COVID-19, therapeutic strategies capable of acting directly on viral gene expression remain relevant both for SARS-CoV-2 and for the broader development of adaptable antiviral platforms against emerging RNA viruses (Maziec et al., 2025).

Accordingly, several nucleic acid-based therapeutic approaches, including small interfering RNAs (siRNAs), antisense oligonucleotides, ribozymes, aptamers, and other functional nucleic acids, have been investigated for targeting viral genomes or transcripts required for infection (Khanali et al., 2021; Tarn et al., 2021). Sequence-specific gene-silencing strategies are particularly attractive because they can, in principle, be redirected toward defined regions of a viral genome. However, several factors constrain their therapeutic application, including limited nuclease stability, inefficient cellular uptake, unfavorable electrostatic interactions with negatively charged cell membranes, and endosomal sequestration (Anwar et al., 2023). In addition, some RNA-based silencing strategies depend on recruitment of endogenous cellular machinery to mediate target degradation (Li & Rana, 2012). These limitations have encouraged the development of alternative nucleic acid-based systems that can directly recognize and catalytically cleave target RNA.

DNAzymes are single-stranded catalytic DNA molecules that cleave target RNA sequences with high sequence specificity, making them promising candidates for gene therapy applications (Thomas et al., 2021; Xiao et al., 2023). Nevertheless, despite their potent catalytic activity, DNAzymes face major barriers to therapeutic translation. They are prone to nuclease-mediated degradation (Xiao et al., 2023), exhibit low stability under physiological conditions, and, most critically, display inefficient cellular uptake, partly due to endosomal entrapment (Yan et al., 2026), the endosomal environment’s low pH value (Qiu et al., 2023), and unfavorable electrostatic interactions with negatively charged cell membranes (Lee et al., 2026). To overcome these obstacles, chemical modification strategies and nanocarrier-based delivery systems, such as lipid, polymer, and inorganic nanoparticles, as well as DNA nanostructures, have been widely explored. While these approaches improve serum stability and cellular uptake, they often come with drawbacks, including toxic side effects, reduced catalytic efficiency, and limited *in vivo* validation (Huo et al., 2020).

Aptamers, on the other hand, have emerged as an appealing alternative for the targeted delivery of nucleic acid therapeutics. Aptamers are short single-stranded DNA or RNA oligonucleotides that fold into distinct three-dimensional structures, enabling them to recognize and bind disease-specific molecular targets with high affinity and specificity (Sun et al., 2014; Wang, 2020). Unlike conventional nanocarriers, aptamers are biocompatible, do not elicit toxic side effects, and can be produced synthetically with high reproducibility (Chandola & Neerathilingam, 2019; H. Yu et al., 2025). Moreover, they have already been used to facilitate the delivery of siRNA, shRNA, and antisense oligonucleotides (ASOs), suggesting their potential as efficient carriers for DNAzymes as well (Kruspe & Giangrande, 2017; Thiel & Giangrande, 2010).

Among the reported aptamers, AS1411 has drawn particular attention. AS1411 is a 26-base guanine-rich DNA aptamer that folds into a G-quadruplex structure with the sequence of 5′-GGTGGTGGTGGTTGTGGTGGTGGTGG-3′, conferring high stability against nucleases and enabling selective binding to nucleolin (Ogloblina et al., 2020), a protein overexpressed on the surface of many cancer and virus-infected cells (Tonello et al., 2022). This unique interaction underlies its ability to accumulate in malignant cells while sparing healthy counterparts preferentially. Beyond its intrinsic antiproliferative activity, AS1411 has advanced to phase II clinical trials in acute myeloid leukemia and renal cell carcinoma, demonstrating both safety and therapeutic potential (Rosenberg et al., 2014; Stuart et al., 2009). In addition, AS1411 has been explored as a versatile delivery module: it can be conjugated to nanoparticles, drugs, or imaging probes, thereby enhancing target specificity and intracellular uptake (Hwang et al., 2010; Verma et al., 2025). AS1411, best known for its applications in cancer therapy due to its high-affinity binding to nucleolin, may also have broader relevance in viral contexts. Nucleolin is a well-established entry factor for several RNA viruses, including HIV-1, Influenza A (Tonello et al., 2022), and respiratory syncytial virus (RSV) (Tayyari et al., 2011). Although direct evidence for SARS-CoV-2 remains limited, the recurring involvement of nucleolin in viral infection pathways provides a plausible rationale for exploring AS1411 in coronavirus-related settings. Moreover, recent studies have described a bidirectional link between COVID-19 and cancer, with infection not only worsening prognosis in cancer patients but also increasing susceptibility to malignancies, particularly in the lung (Dai et al., 2020; Kwan et al., 2021; Wallace et al., 2025). This dual evidence provides a biologically grounded justification for investigating AS1411-mediated delivery in pulmonary models relevant to both oncological and viral applications.

For such a strategy, the Spike (S) gene represents a particularly relevant target because its encoded glycoprotein is essential for host-cell recognition, membrane fusion, and viral entry (S. Yu et al., 2022). Therefore, sequence-specific cleavage of the S transcript offers a direct approach to suppressing expression of a key viral protein. In addition, relatively conserved regions have been identified within the Spike protein, including regions of the S2 subunit involved in the membrane-fusion machinery (Polo-Megías et al., 2022). Such conservation provides a rational basis for considering these regions as targets for sequence-directed catalytic oligonucleotides. However, because the accessibility and structural context of an RNA target can substantially influence DNAzyme activity (Cairns et al., 1999; Nurmi et al., 2024), identification of an effective Spike-targeting DNAzyme requires comparative experimental evaluation rather than reliance on catalytic-core identity alone.

Integrating these considerations, the combination of an efficient Spike RNA-cleaving DNAzyme with AS1411 was investigated as a strategy to incorporate catalytic RNA cleavage and cellular targeting within a single molecular construct. In this study, a panel of DNAzymes representing different catalytic cores and target regions within the SARS-CoV-2 Spike RNA was designed and comparatively evaluated for in vitro cleavage activity. Following identification of the lead catalytic candidate, the DNAzyme was conjugated to AS1411 in two opposite linear orientations to generate aptamer–DNAzyme chimeras, which were subsequently evaluated in a cellular model expressing the Spike gene. Because conjugation may alter the structural organization and accessibility of either functional module, the potential influence of orientation was further investigated using circular dichroism spectroscopy, three-dimensional structural modeling, RNA–complex analysis, and molecular docking with nucleolin. Through this integrated approach, a design strategy combining catalytic activity, cellular applicability, and molecular organization within a single aptamer–DNAzyme platform targeting SARS-CoV-2 Spike RNA was evaluated.

## MATERIALS AND METHODS

### Viral RNA Production from the *S Gene*

Viral RNA was synthesized *in vitro* using T7 polymerase following the manufacturer’s protocol (Thermo Fisher Scientific). The reaction mixture contained template DNA (synthetically obtained *S gene* with a 5′ T7 promoter, see supporting information), transcription buffer, NTP mix, T7 polymerase, RiboLock RNase inhibitor, DTT, and nuclease-free water. The samples were incubated at 37°C for 16 hours. The transcription products were visualized on a 1% agarose gel, confirming successful RNA synthesis at the expected yield. Unless otherwise stated, all electrophoreses are utilized with SafeView dye.

### *In Vitro* DNAzyme Catalytic Activity Assay

To evaluate the catalytic activity of the DNAzymes on the synthesized viral RNA, a panel of sequences designed using the DNAzymeBuilder bioinformatics tool (10-23-1076, 10-23-624, 8-17-531, 8-17-769, AA14-818, AA14-931, AC17-819, and AA17-932) were purchased from Sentebiolab, Ankara. Before initiating the time-dependent cleavage assays, a preliminary dose-response experiment was performed with DNAzyme 8-17-769 **(ref)** to determine the optimal working concentration. Viral RNA was diluted in MgCl₂ buffer, and reaction mixtures were prepared using target RNA at a final concentration of 60 nM and serial dilutions of DNAzyme 8-17-769, resulting in final DNAzyme concentrations of 20 µM, 4 µM, 0.8 µM, 0.32 µM, and 0.16 µM. The reactions were incubated overnight at 37 °C. Following incubation, 60% formamide was added to prevent secondary folding of both RNA and DNAzymes, and the samples were heated at 65 °C, right before agarose gel electrophoresis. Reaction products were visualized by 1% agarose gel electrophoresis.

Cleavage assays were performed for all DNAzymes in a time-dependent manner. For each reaction, viral RNA diluted in MgCl₂ buffer was combined with DNAzyme at a final concentration of 20 µM. The reactions were incubated at 37 °C with agitation at 400 rpm for 24 h, 48 h, or 72 h as indicated. Control reactions lacking DNAzyme were included. During incubation, the mixtures were sampled at various time steps. After incubation, 60% formamide was added to each sample, followed by heating at 65 °C, right before agarose gel electrophoresis to renature the structures. Cleavage products were analyzed on 1% agarose gels to assess the catalytic activity of each DNAzyme over time.

### Band Quantification and Kinetic Calculations

Gel images from RNA cleavage assays were quantified using ImageJ. For each lane, vertical intensity profiles were generated, extending from the intact RNA band to the lowest detectable cleavage product. The line width was kept constant within each gel to ensure comparable lane sampling. Intensity profiles obtained for each time point were exported for further analysis.

To normalize lane profiles, the initial intact RNA signal (*Y*_0_) and the endpoint cleavage signal (*Y*_∞_) were used as reference profiles. The theoretical intensity (*Y_t_*) profile at each time point was calculated according to eqn. (X):

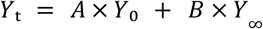

where A and B are normalization coefficients. These coefficients were optimized by minimizing the sum of squared differences between the observed and theoretical intensity profiles. The optimized coefficients were then used to calculate the fractional amount of uncleaved substrate remaining at each time point, *f*(*t*). The observed substrate concentration was calculated using eqn. (X):

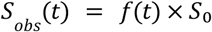

where *S_obs_* (*t*) represents the amount of uncleaved RNA remaining at time *t*, and *S*₀ is the initial substrate concentration of 60 nM.

Cleavage kinetics were analyzed under a pseudo-first-order model, based on the excess of DNAzyme relative to RNA substrate. The decrease in uncleaved RNA over time was described by eqn. (X):

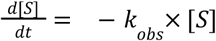

After integration, substrate decay over time was expressed as eqn. (X):

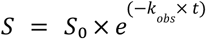

For kinetic fitting, ln-transformed substrate concentrations were fitted to the linearized pseudo-first-order model shown in eqn. (X):

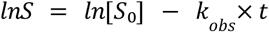

For each time point, the calculated substrate concentration was obtained from the model as shown in eqn. (X):

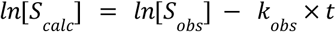

The observed and calculated *ln*[*S*] values across all time points were compared, and *k_obs_* was optimized by minimizing the sum of squared residuals using the GRG Nonlinear method. The resulting *k_obs_* values were used to compare the cleavage rates of different DNAzymes, with higher *k_obs_* values indicating faster substrate degradation and greater apparent catalytic activity.

For graphical representation of cleavage kinetics, normalized fraction values were plotted as a function of reaction time using GraphPad Prism. Pseudo-first-order decay curves were generated using the optimized *k_obs_* values obtained from the kinetic analysis and the exponential decay relationship described above. These fitted curves were overlaid with the experimentally derived fraction values to visualize the time-dependent cleavage profiles of each DNAzyme.

### Circular Dichroism Spectrometry

The structural properties of the aptamer, the DNAzyme, and their conjugated forms were investigated using circular dichroism (CD) spectroscopy. Oligonucleotide sequences corresponding to 8-17-769, AS1411, 8-17-769–AS1411, and AS1411–8-17-769 were prepared at a final concentration of 10 µM in a buffer containing 10 mM Tris and 100 µM KCl. CD measurements were performed using a CD spectrometer over the wavelength range of 190–400 nm, with three replicate scans acquired for each sample. CD, voltage, and absorbance spectra were obtained, and the alignment of the observed peaks was compared with characteristic peak positions reported for known G-quadruplex structures to evaluate G-quadruplex formation and structural integrity.

### Creating an A549 cell line model expressing the *S gene*

To establish a mammalian cell–based gene expression model, a pTwist_cDNA_EF1α plasmid (see supplementary information for plasmid map) encoding the SARS-CoV-2 *Spike (S) gene* and Woodchuck hepatitis virus posttranscriptional regulatory element (WPRE), which was purchased from Sentromer/ TwistBio, was used. The plasmid, which was designed for high-level gene expression in mammalian cells, contains the EF1α promoter and a neomycin resistance gene. Plasmid stocks were amplified in *Escherichia coli* DH5α cells via heat-shock transformation. Briefly, competent cells were thawed on ice and then mixed with plasmid DNA and incubated on ice. Cells were heat-shocked at 42 °C and immediately transferred to ice. Then they recovered in S.O.C. medium at 37 °C with shaking. Transformed cells were plated on LB agar containing ampicillin and incubated overnight at 37 °C. Ampicillin-resistant colonies were selected, expanded in LB medium, and plasmid DNA was isolated using the EZ-10 plasmid isolation kit.

AS1411–8-17-769 and 8-17-769–AS1411 conjugates were synthetically designed with the sequence of “AS1411_8-17-769: 5’-GGTGGTGGTGGTTGTGGTGGTGGTGGTTTTGCTGTCCAATTCCAGCGGATCGAATGAAGAAGAA-3’, 8-17-769_AS1411: 5’-GCTGTCCAATTCCAGCGGATCGAATGAAGAAGAATTTTGGTGGTGGTGGTTGTGGTGGTGGTGG-3’” and obtained (from Sentebiolab). The conjugates were engineered such that the aptamer and DNAzyme were connected via a four-thymine (T) linker, providing structural flexibility. Thymine residues were selected to avoid potential interference with G-quadruplex formation, which could occur with cytosine-containing linkers.

A549 cells were cultured in complete RPMI medium and used to establish the gene expression model. Initial transfection experiments were performed using TurboFectin 8.0. Cells were seeded into 6-well plates at a density of 2×10^5^ cells and transfected on the following day (after 24 h incubation) at 50–70% confluency. Transfection complexes were prepared by diluting of plasmid DNA in Opti-MEM and mixing it with TurboFectin transfection agent, followed by a 15-minute incubation. Complexes were added to each well, and cells were incubated for 48 h. Cells were then washed with PBS, detached using trypsin, and collected by centrifugation. Total RNA was isolated using the Monarch RNA isolation kit, and cDNA synthesis was performed using the a.b.m. one script plus cDNA synthesis kit. Since the expression modeling strategy does not involve permanent integration of the gene fragment into the cellular genome, random primers were used for cDNA synthesis.

Quantitative PCR (qPCR) targeted the *S gene*, with the WPRE used as an internal reference. Naked plasmid DNA served as a positive control. qPCR reactions were prepared using iTaq SYBR Supermix.

### Knockdown of the *S gene* in the A549 cell line models via DNAzyme and its conjugates

A second set of transfection experiments was conducted using the Xfect transfection reagent. Cells were seeded into 6-well plates at a density of 2×10^5^ cells and transfected on the following day (after 24 h incubation) at 50–70% confluency, using plasmid DNA mixed with Xfect polymer and buffer according to the manufacturer’s instructions (TAKARA-bio). After incubation, nanoparticle complexes were added to each well. The medium was replaced after 4 h, and cells were incubated for an additional 48 h.

To induce gene silencing, DNAzyme 8-17-769 was delivered into cells 48 hours after plasmid transfection using Xfect-based nanoparticle complexes, at the peak point of expression. In parallel, AS1411–8-17-769 and 8-17-769–AS1411 conjugates, each diluted separately in 1X TE buffer, were added to additional wells to evaluate aptamer-mediated delivery efficiency. A reference well expressing the *S gene* without any silencing molecule was included. Throughout the experiment, cells were maintained under selective conditions using G418/neomycin 48 hours after the addition of silencing molecules.

Total RNA was isolated using the Monarch RNA isolation kit, and cDNA synthesis was performed using random primers and the a.b.m. one script plus cDNA synthesis kit. Quantitative PCR (qPCR) targeted the *S gene*, with the WPRE used as an internal reference. Naked plasmid DNA served as a positive control. qPCR reactions were prepared using iTaq SYBR Supermix.

qPCR data were analyzed using Cq values exported from the CFX Maestro software. Technical replicates were averaged after excluding inconsistent outliers. Relative *S gene* expression was calculated using the Cq method (2^−ΔΔC^*_q_* method/ Livak), with WPRE used as the internal reference and the untreated *S gene-expressing* cells used as the negative control. Relative fold change values were calculated as 2^−ΔΔC^*_q_*. Statistical significance between treatment groups and the control group was assessed using two-tailed t-tests on ΔCq values, and p < 0.05 was considered statistically significant.

## RESULTS AND DISCUSSION

### In *Vitro* Catalytic Characterization of DNAzymes Targeting SARS-CoV-2 Spike RNA and Identification of the Lead Candidate

To identify candidate deoxyribozymes capable of exhibiting catalytic activity against SARS-CoV-2 Spike RNA, a total of eight deoxyribozymes representing different catalytic cores and target regions were designed. The resulting panel comprised sequences containing the 8-17, 10-23, AA14, AC17, and AA17 catalytic motifs, each directed against distinct positions within the Spike RNA (Table 1). Rather than evaluating the activity of a single catalytic core in isolation, this design enabled a comparative assessment of different catalytic core–target site combinations against the same RNA substrate.

**Table 1.**
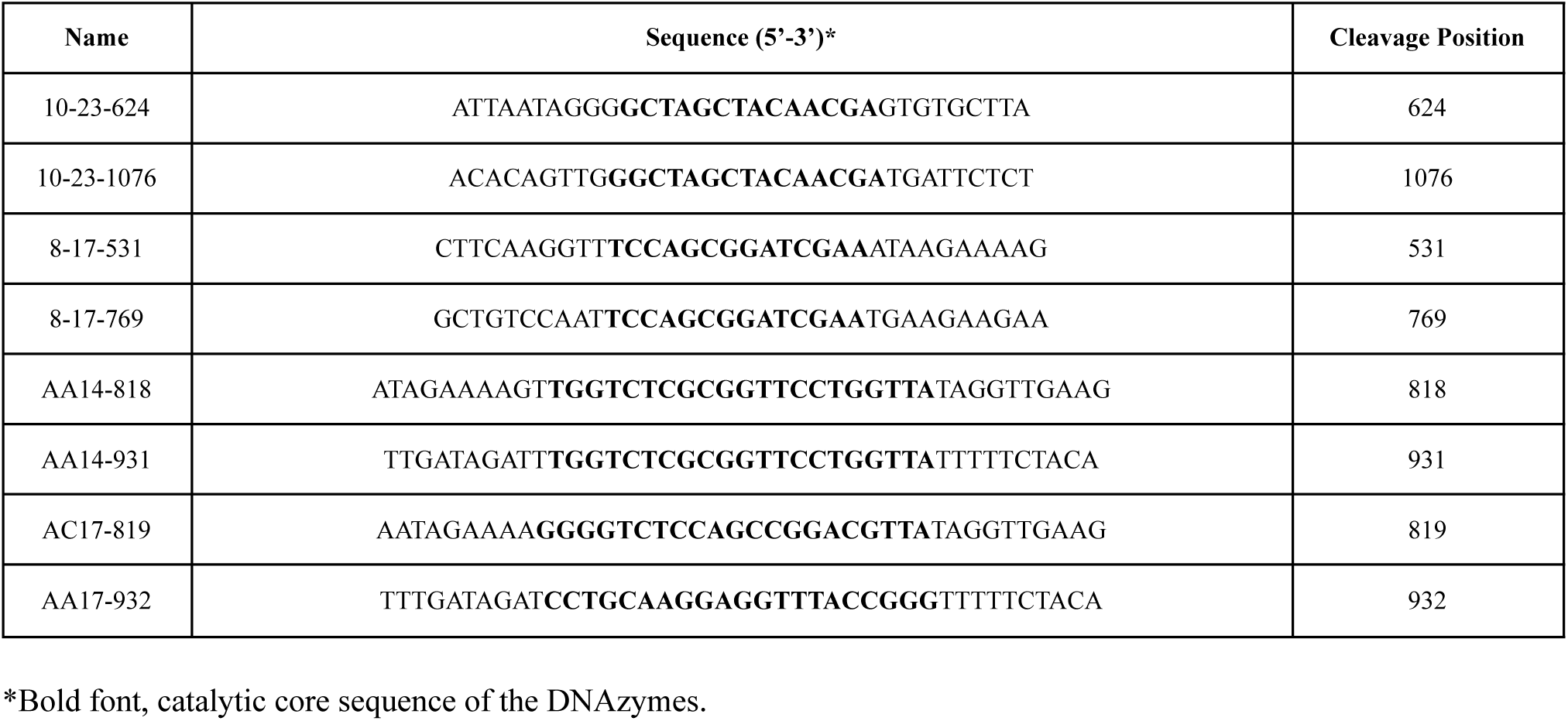
The sequences of eight DNAzymes targeting the Spike gene mRNA.

To enable comparison of the time-dependent cleavage profiles of the candidates, the reaction conditions were first standardized through a preliminary concentration screening using 8-17-769. Partial substrate loss was observed at lower deoxyribozyme concentrations, whereas pronounced and near-complete RNA cleavage was achieved by the end of the incubation period in the presence of 20 µM deoxyribozyme (Figure 1A). Therefore, comparative time-dependent cleavage experiments among the candidates were conducted at a deoxyribozyme concentration of 20 µM. The purpose of this approach was not to define a therapeutically optimal dose, but rather to establish a common experimental condition under which the relative cleavage performances of the tested sequences could be compared.

**Figure 1.**
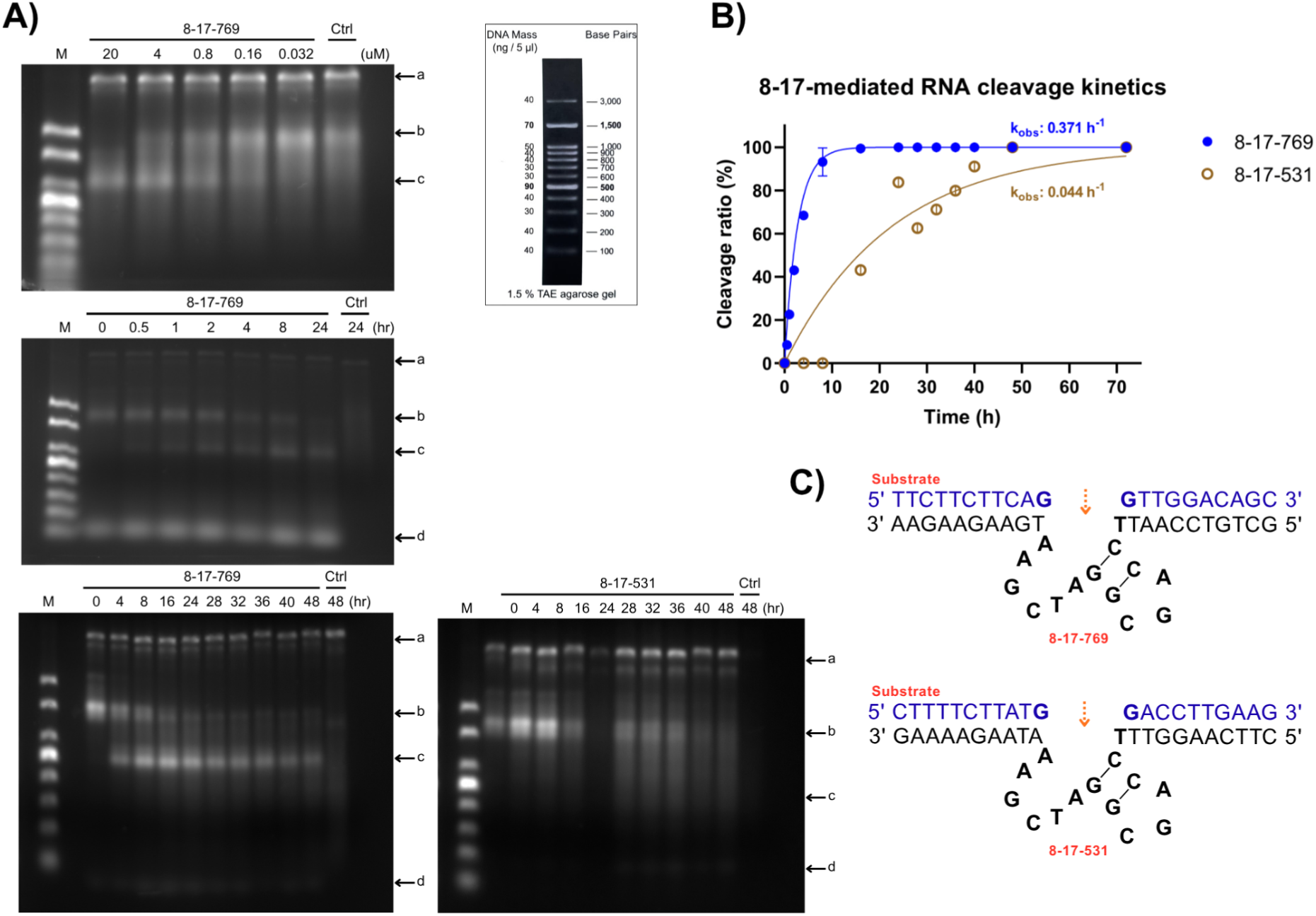
A) Agarose gel electrophoresis results of cleavage activity assays performed for the designed 8-17 DNAzyme variants (8-17-769, 8-17-531). Representative results from experiments conducted over 24 h and extended to 48 h incubation periods are shown. In addition, the results of a preliminary dose–response assay performed with DNAzyme 8–17–769 are presented (top left). A DNA size marker, whose band positions are indicated in the reference map, was used in all gels. The bands observed at the indicated positions correspond to the following molecules: **a)** template DNA (*S gene*), **b)** uncleaved *S gene* viral RNA (substrate), **c)** cleaved *S gene* viral RNA (product), **d)** DNAzyme. **B)** Cleavage kinetics of 8–17–769 and 8–17–531 DNAzymes based on quantified gel intensities. Data points indicate experimentally derived cleavage ratios, and fitted curves represent pseudo-first-order kinetic profiles generated using the calculated *k_obs_* values. **C)** Schematic representation of the predicted substrate-binding configurations of the 8–17–769 and 8–17–531 DNAzymes. The target RNA substrate sequences are shown in blue, while the DNAzyme binding arms and catalytic core are shown in black. The orange arrows indicate the predicted RNA cleavage sites.

Under these conditions, time-course experiments revealed marked target-site-dependent differences in performance, even among deoxyribozymes sharing the same catalytic motif. Among the 8-17 variants, 8-17-769 showed pronounced substrate loss from the early time points and eliminated most of the uncleaved RNA within the first 24 h; a similar cleavage profile observed in the extended time-course experiment supported the reproducibility of this activity (Figure 1A). In contrast, 8-17-531, which contains the same 8-17 catalytic core but targets a different region of the Spike RNA, did not produce a distinct cleavage product band. Instead, an increase in signal dispersion within the product region, together with a gradual decrease in substrate band intensity from approximately 4 h onward, was consistent with a slower cleavage process; even after 48 h of incubation, cleavage remained lower than that observed for 8-17-769 (Figure 1A).

Quantitative analysis of the intensity profiles derived from the gel images further clarified this distinction. While 8-17-769 exhibited an early and rapid decrease in substrate abundance, 8-17-531 displayed a slower time-dependent profile (Figure 1B). Despite sharing the same catalytic core, the markedly different behaviors of the two variants at distinct target sites indicate that the observed activity cannot be explained solely by the identity of the catalytic motif. The local sequence context of the target site and/or RNA accessibility may contribute to this difference (Nakajima et al., 2026); however, because these parameters were not independently evaluated in the present study, the observed difference cannot be directly attributed to a specific structural mechanism. The predicted substrate–deoxyribozyme binding configurations of the two variants are shown in Figure 1C.

A similar target-site dependence was also observed among the 10-23 variants. Although no distinct product band was detected for 10-23-624 at the early time points, a decrease in substrate band intensity accompanied by an increase in signal within the product region became evident from approximately 16 h onward, consistent with a slow but measurable cleavage activity (Figure 2A). In contrast, no meaningful substrate loss or cleavage product was detected for 10-23-1076 over the 48 h incubation period, and this candidate was therefore excluded from subsequent quantitative kinetic analysis. Time-dependent profiles calculated from gel intensities likewise supported the low-level activity of 10-23-624, whereas no measurable cleavage profile could be obtained for 10-23-1076 (Figure 2B). These findings further demonstrated that directing the same 10-23 catalytic core to different regions of the Spike RNA did not result in functionally equivalent cleavage performance (Figure 2C).

**Figure 2.**
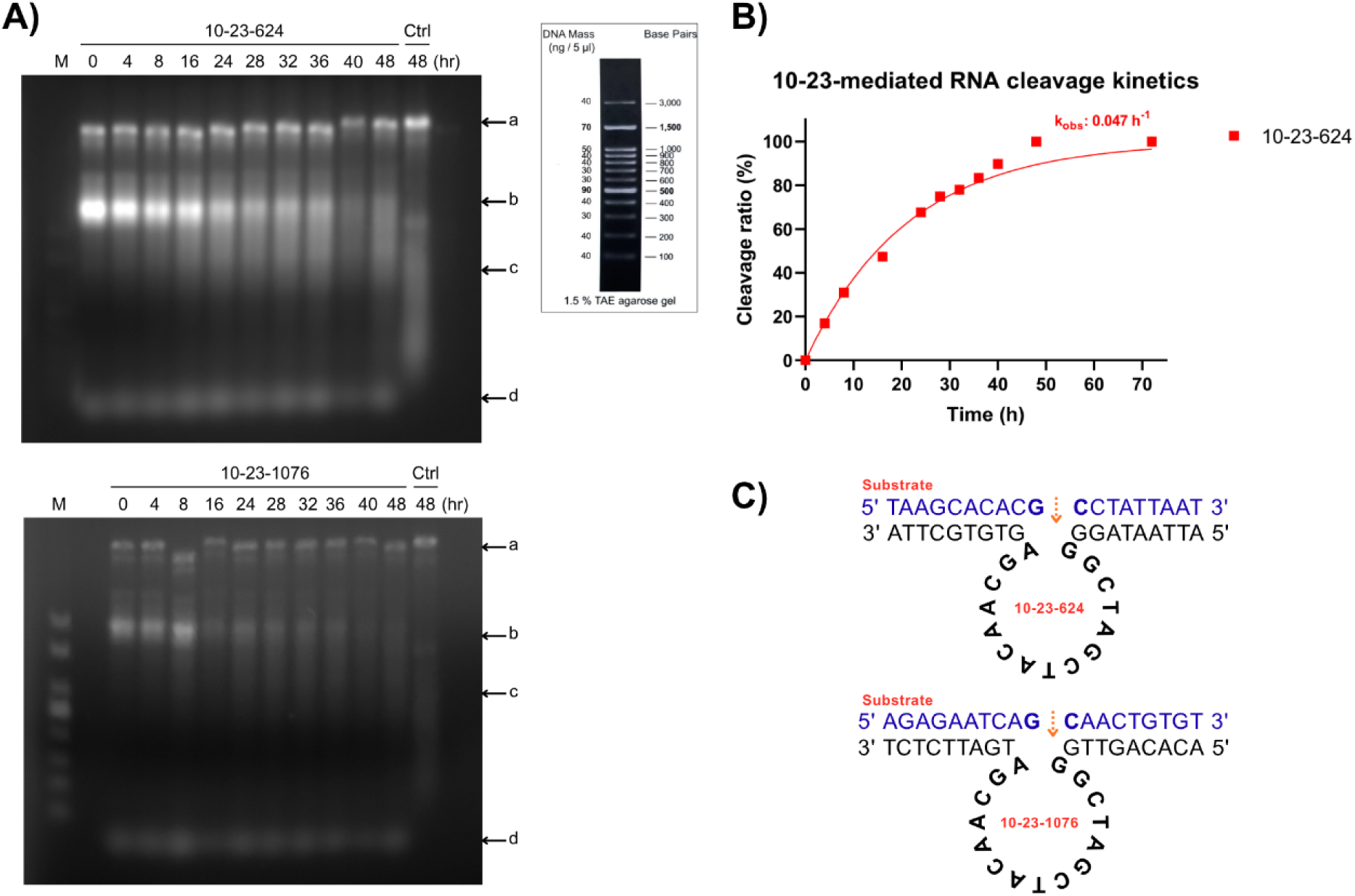
Agarose gel electrophoresis results of cleavage activity assays performed for the designed 10–23 DNAzyme variants (10-23-624, 10-23-1076). Representative results from experiments conducted over 24 h and extended to 48 h incubation periods are shown. A DNA size marker, whose band positions are indicated in the reference map, was used in all gels. The bands observed at the indicated positions correspond to the following molecules: **a)** template DNA (*S gene*), **b)** uncleaved *S gene* viral RNA (substrate), **c)** cleaved *S gene* viral RNA (product), **d)** DNAzyme. **B)** Cleavage kinetics of 10-23-624 and 10-23-1076 DNAzymes based on quantified gel intensities. Data points indicate experimentally derived cleavage ratios, and fitted curves represent pseudo-first-order kinetic profiles generated using the calculated *k_obs_* values. **C)** Schematic representation of the predicted substrate-binding configurations of the 10-23-624 and 10-23-1076 DNAzymes. The target RNA substrate sequences are shown in blue, while the DNAzyme binding arms and catalytic core are shown in black. The orange arrows indicate the predicted RNA cleavage sites.

Catalytic activity was observed for both AA14 variants, although their cleavage profiles progressed more slowly than that of 8-17-769. Substrate loss became evident from approximately 8 h onward for both AA14-818 and AA14-931, with cleavage reaching completion at around 48 h for AA14-818 and at later time points for AA14-931 (Figure 3A). This slower kinetic behavior was also reflected in the time-dependent cleavage profiles derived from gel intensity measurements (Figure 3B). Thus, although both AA14 variants were capable of cleaving Spike RNA, their performance at early time points was lower than that of the more rapidly acting candidates.

**Figure 3.**
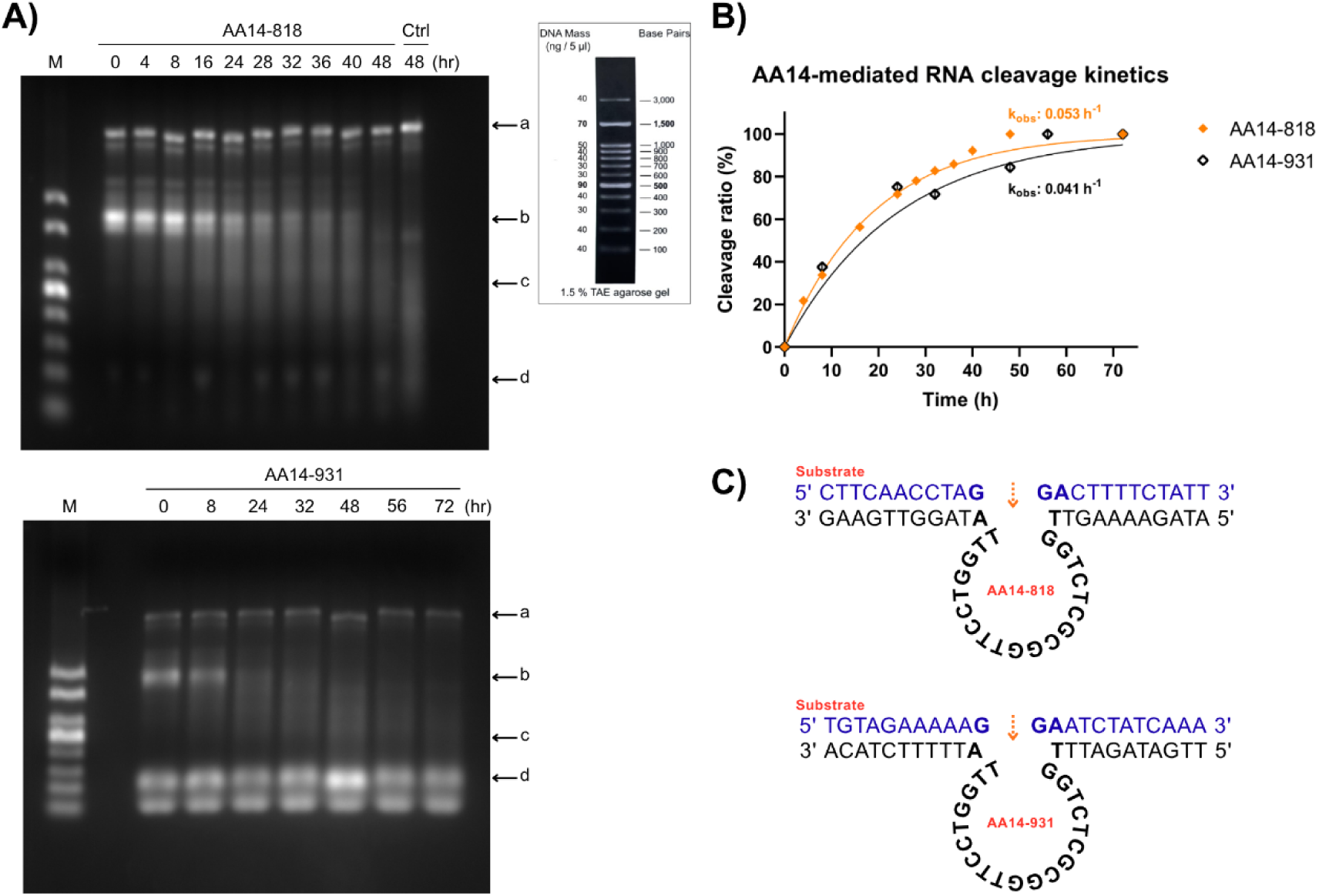
Agarose gel electrophoresis results of cleavage activity assays performed for the designed AA14 DNAzyme variants (AA14-818, AA14-931). Representative results from experiments conducted over 24 h and extended to 48 h incubation periods are shown. A DNA size marker, whose band positions are indicated in the reference map, was used in all gels. The bands observed at the indicated positions correspond to the following molecules: **a)** template DNA (*S gene*), **b)** uncleaved *S gene* viral RNA (substrate), **c)** cleaved *S gene* viral RNA (product), **d)** DNAzyme. **B)** Cleavage kinetics of AA14-818 and AA14-931 DNAzymes based on quantified gel intensities. Data points indicate experimentally derived cleavage ratios, and fitted curves represent pseudo-first-order kinetic profiles generated using the calculated *k_obs_* values. **C)** Schematic representation of the predicted substrate-binding configurations of the AA14-818 and AA14-931 DNAzymes. The target RNA substrate sequences are shown in blue, while the DNAzyme binding arms and catalytic core are shown in black. The orange arrows indicate the predicted RNA cleavage sites.

Distinct yet divergent cleavage behaviors were also observed among the variants carrying the AA17 motif. AC17-819 was one of the candidates showing early catalytic activity, with cleavage becoming evident within the first hour and progressing to substantial substrate loss within the first 24 h (Figure 4A). In contrast, cleavage by AA17-932 began later, with clear activity emerging after approximately 8 h and near-complete substrate loss requiring a longer incubation period. This difference was likewise reflected in the quantitative time-dependent profiles (Figure 4B).

**Figure 4.**
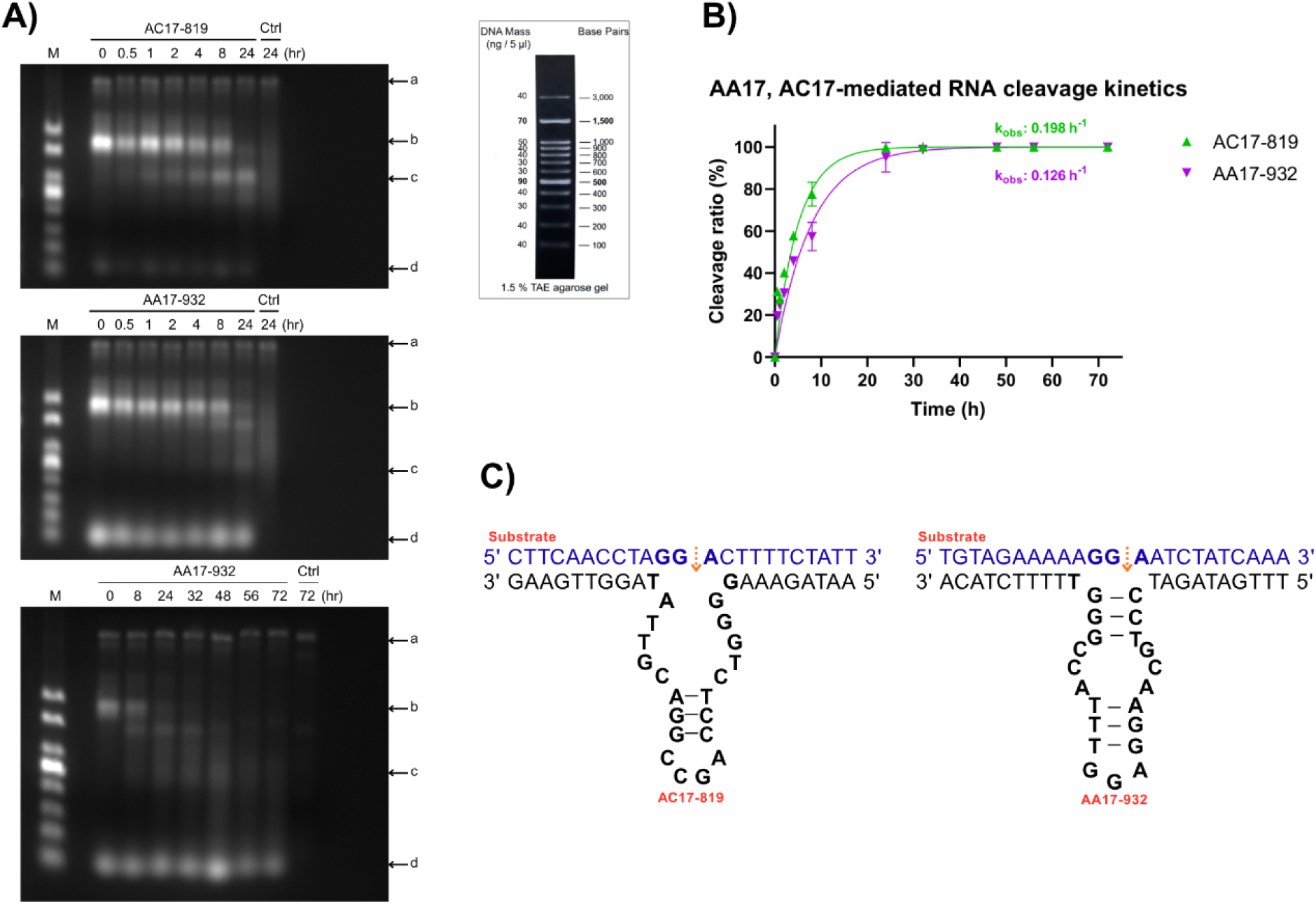
Agarose gel electrophoresis results of cleavage activity assays performed for the designed AA17 DNAzyme variants (AC17-819, AA17-932). Representative results from experiments conducted over 24 h and extended to 48 h incubation periods are shown. A DNA size marker, whose band positions are indicated in the reference map, was used in all gels. The bands observed at the indicated positions correspond to the following molecules: **a)** template DNA (*S gene*), **b)** uncleaved *S gene* viral RNA (substrate), **c)** cleaved *S gene* viral RNA (product), **d)** DNAzyme. **B)** Cleavage kinetics of AC17-819 and AA17-932 DNAzymes based on quantified gel intensities. Data points indicate experimentally derived cleavage ratios, and fitted curves represent pseudo-first-order kinetic profiles generated using the calculated *k_obs_* values. **C)** Schematic representation of the predicted substrate-binding configurations of the AC17-819 and AA17-932 DNAzymes. The target RNA substrate sequences are shown in blue, while the DNAzyme binding arms and catalytic core are shown in black. The orange arrows indicate the predicted RNA cleavage sites.

To quantitatively compare the gel-based observations across candidates, time-dependent substrate intensities were analyzed using ImageJ, and the observed rate constants *k_obs_* were calculated according to a pseudo-first-order model (Table 2). This analysis revealed a clear performance ranking that was consistent with the qualitative gel findings. Among the tested candidates, 8-17-769 exhibited the highest observed cleavage rates, with *k_obs_* values of 0.0071 and 0.0052 in two independent time-course experiments, respectively. This was followed by AC17-819 (0.0030 and 0.0036) and AA17-932 (0.00221 and 0.0020). Notably, for these three candidates, the close agreement between values obtained from independent experiments performed using different time intervals indicated that their relative performance ranking was maintained across experiments (Table 2).

**Table 2.** Calculated *k_obs_* values derived from fractional cleavage data obtained at each time interval. Values are listed in descending order according to cleavage efficiency. *N/D: Not detected, “-”: Measurement not performed.

| DNAzyme / $k_{obs}$ | 1<sup>st</sup> measurement | 2<sup>nd</sup> measurement |
| --- | --- | --- |
| **8-17-769** | 0.0071 | 0.0052 |
| **AC17-819** | 0.0030 | 0.0036 |
| **AA17-932** | 0.00221 | 0.0020 |
| **AA14-818** | 0.00089 | - |
| **10-23-624** | 0.00078 | - |
| **8-17-531** | 0.00074 | - |
| **AA14-931** | 0.00069 | - |
| **10-23-1076** | N/D | N/D |

In contrast, the *k_obs_* values calculated for AA14-818, 10-23-624, 8-17-531, and AA14-931 were lower, consistent with the delayed substrate loss observed in their gel profiles. For 10-23-1076, no reliable cleavage signal was detected during the experimental period, and *k_obs_* could therefore not be calculated. Taken together, the time-dependent cleavage patterns observed by gel electrophoresis and the quantitative kinetic analysis indicated that the differences in performance among the candidates were determined not only by the extent of cleavage at the endpoint, but also by the rate at which cleavage occurred.

Among all candidates, 8-17-769 was identified as the strongest lead candidate because it exhibited pronounced substrate loss at early time points, achieved the highest *k_obs_* values in two independent experimental series, and showed a reproducible cleavage profile. Accordingly, 8-17-769 was selected as the catalytic component of the chimeric constructs to be generated with the AS1411 aptamer in the subsequent stage of the study to confer a cellular delivery function. Thus, the *in vitro* screening stage not only identified sequences capable of cleaving Spike RNA, but also established a rationally selected catalytic component for the aptamer–deoxyribozyme construct to be evaluated in the cellular system.

### Functional Evaluation of AS1411–8-17-769 Chimeras in a Cellular Spike Expression Model

The observation that 8-17-769 exhibited the fastest and most reproducible Spike RNA cleavage profile among the tested deoxyribozymes in the *in vitro* cleavage assays raised the question of whether this candidate could be translated into a functional silencing agent under cellular conditions. To address this, a cellular model enabling controlled expression of the SARS-CoV-2 Spike gene was first established. The pTwist_cDNA_EF1α plasmid, carrying the Spike gene under the control of the EF1α promoter, was transfected into A549 cells, and qPCR analysis of RNA isolated from the cells detected amplification signals for both the *S gene* and the WPRE region located within the same expression cassette. These results confirmed that the Spike transcript could be expressed in A549 cells for use in subsequent silencing experiments.

Following establishment of the cellular model, the effect of 8-17-769, identified as the lead candidate in the in vitro screening, was comparatively evaluated in a cellular context together with the contribution of AS1411 conjugation to this effect. For this purpose, 8-17-769 was delivered into cells using Xfect, while the AS1411–8-17-769 and 8-17-769–AS1411 chimeras, in which the aptamer and deoxyribozyme modules were connected by a four-thymine linker, were evaluated in separate experimental groups. This design enabled assessment not only of the cellular activity of the deoxyribozyme itself, but also of whether arranging the same two functional modules in different linear orientations influenced silencing performance.

In the qPCR analysis, lower relative S-gene expression levels were observed in the experimental groups containing 8-17-769 compared with the reference group expressing the *S gene* alone (Figure 5). Although free 8-17-769 delivered into the cells using Xfect was also associated with reduced expression, a more pronounced decrease was observed in the AS1411-containing chimeric constructs. According to the relative expression analysis, S-gene expression was reduced by approximately fivefold in the AS1411–8-17-769 conjugate relative to the reference group, whereas the 8-17-769–AS1411 conjugate exhibited the strongest silencing profile among the tested constructs, with an approximately tenfold reduction (Figure 5). In the statistical analysis reported in the study, significance was observed only for the 8-17-769–AS1411 group (p < 0.05).

**Figure 5.**
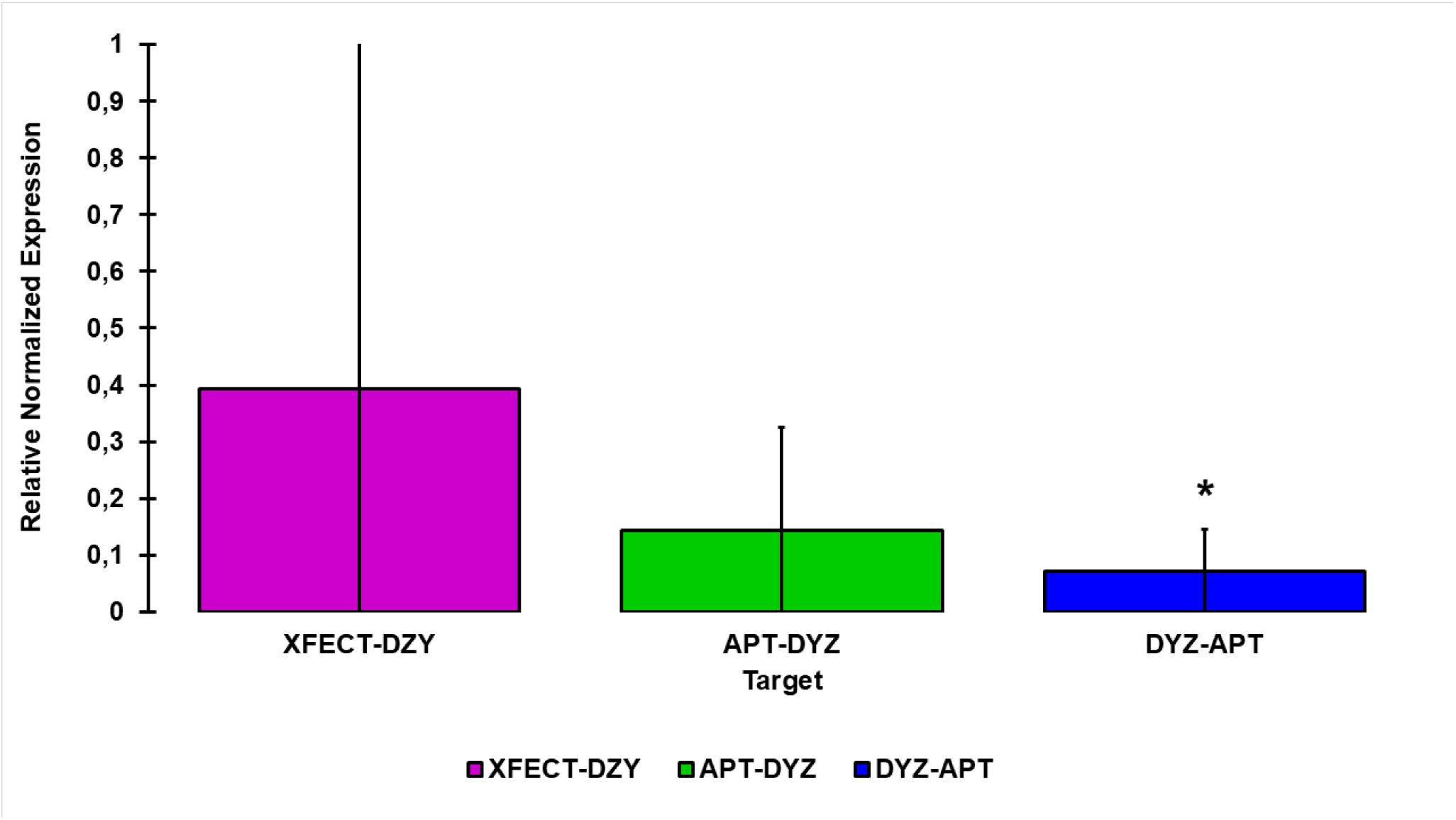
Box plot representation of qPCR results obtained after knock-down experiments. Relative normalized expression levels of the Spike (S) gene are shown for three experimental groups (APT–DYZ, DYZ–APT, and XFECT: DYZ) (DYZ: 8-17-769; APT: AS1411; XFECT: lipid-based transfection reagent). A marked reduction in *S gene* expression was observed in all groups containing 8-17-769, with the most pronounced decrease detected in the DYZ–APT group. **\****p < 0.05*.

This finding is important for two reasons. First, the observation that the RNA-cleaving capacity of 8-17-769 demonstrated *in vitro* was accompanied by reduced S-gene expression in the cellular expression model suggests that the activity of this candidate was not limited to an isolated RNA substrate. Second, and more notably, the nonequivalent performance of the two chimeras containing the same AS1411 and 8-17-769 modules suggests that conjugation orientation is not functionally neutral. The stronger reduction in expression associated with the 8-17-769–AS1411 configuration, in which 8-17-769 is positioned at the 5′ end and AS1411 at the 3′ end, raises the possibility that the intramolecular organization of the two modules may influence the accessibility of either the aptamer or the catalytic region.

However, the available cellular data are insufficient to determine the mechanism underlying this difference in performance. In particular, the qPCR data do not allow discrimination between whether the stronger effect of 8-17-769–AS1411 results from greater cellular uptake, improved accessibility of the deoxyribozyme to the target RNA following cellular entry, or differential effects of conjugation on the structural integrity of the two functional modules. Therefore, the potential basis of the functional difference between the two chimeras was subsequently evaluated at the structural and molecular levels.

### Structural and In Silico Analysis of the Effects of Conjugation Orientation on Chimera Architecture and Molecular Interactions

Although the 8-17-769–AS1411 and AS1411–8-17-769 chimeras contained the same functional components, their distinct silencing profiles in the cellular experiments prompted investigation of the effect of conjugation orientation on molecular structure and target interactions. In particular, the more pronounced reduction in S-gene expression associated with 8-17-769–AS1411 suggested that functional performance may depend not only on the presence of the aptamer and deoxyribozyme modules within the same molecule, but also on their linear organization. Accordingly, the two conjugates were first compared experimentally using circular dichroism (CD) spectroscopy, after which potential structural differences were further evaluated through three-dimensional modeling, RNA–complex analysis, and molecular docking studies with nucleolin.

In the CD spectroscopy analysis, the individual spectral profiles of 8-17-769 and AS1411 were first determined, and the normalized cumulative signal of these profiles was compared with the spectra obtained from the two chimeras (Figure 6A–B). The spectrum of 8-17-769–AS1411 largely overlapped with the cumulative reference profile of the individual components, whereas a more pronounced deviation from the reference profile was observed for the AS1411–8-17-769 conjugate (Figure 6B). These findings indicate that, in the 8-17-769–AS1411 orientation, the two functional modules relatively better preserved their original structural characteristics following conjugation, whereas the reverse orientation was associated with a more pronounced perturbation of the molecular organization.

**Figure 6.**
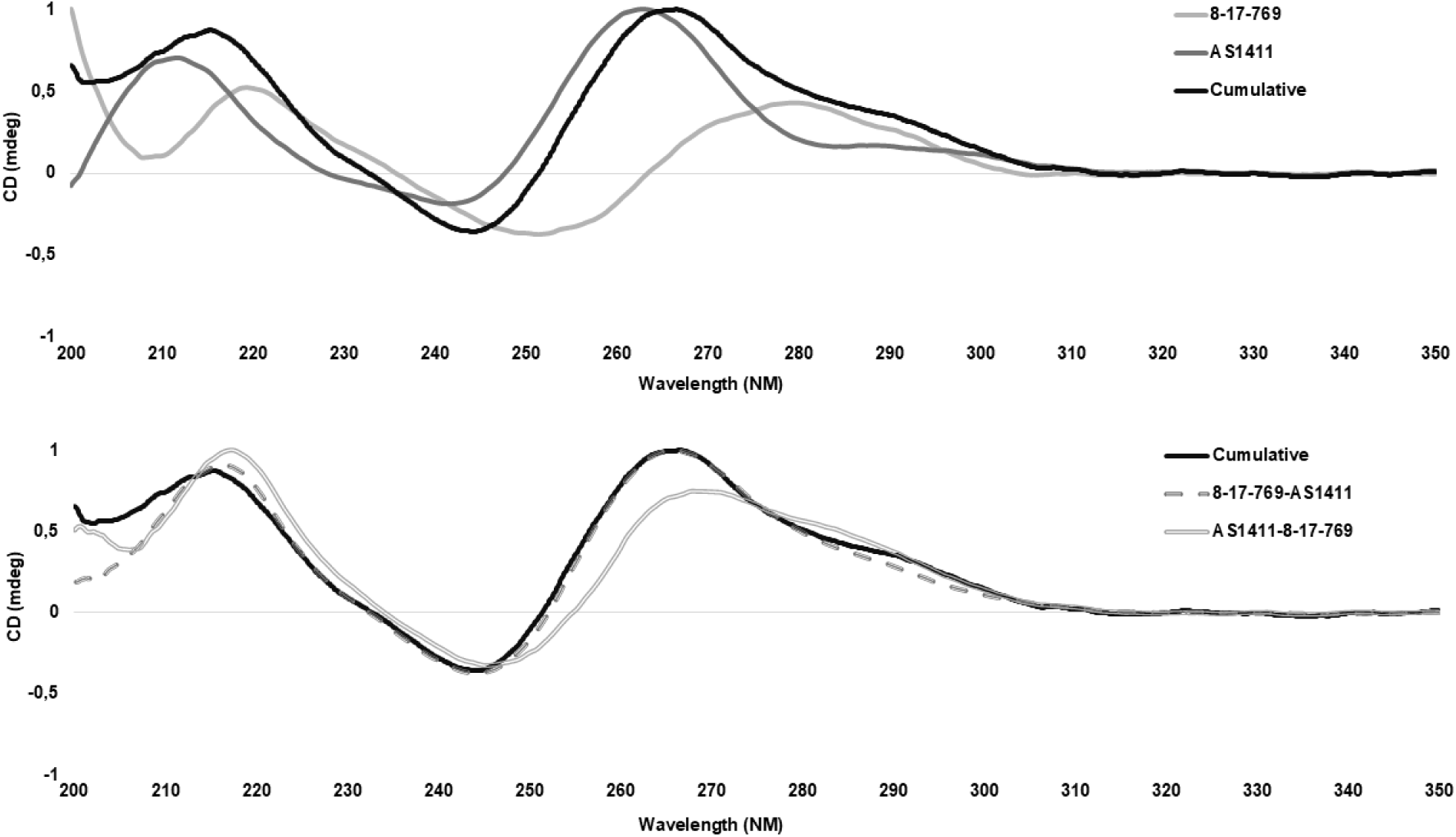
Circular dichroism (CD) spectra obtained from CD spectroscopy measurements. **a)** Overlay of the CD wavelength spectra of 8-17-769, AS1411, and their conjugated forms, normalized to the 0–1 range to allow direct comparison of spectral profiles. **b)** Comparison of the CD spectra of 8-17-769–AS1411 and AS1411–8-17-769 with the cumulative reference curve. The alignment of the 8-17-769–AS1411 spectrum with the cumulative curve indicates that conjugation does not induce substantial structural alterations.

This structural difference is particularly noteworthy when considered alongside the preceding cellular results. The 8-17-769–AS1411 chimera, which exhibited the greater silencing effect, also showed a CD profile that more closely resembled the cumulative behavior of the individual components, whereas AS1411–8-17-769, which displayed the lower silencing profile, exhibited more pronounced spectral alterations. Thus, the CD data are consistent with the hypothesis that conjugation orientation may influence the structural integrity of the chimera and that these structural differences may contribute to the observed variation in functional performance. However, because CD spectra alone cannot define a specific three-dimensional conformation or identify which module is primarily affected, computational modeling was subsequently employed to investigate the molecular basis of the differences between the two chimeras in greater detail.

In addition to the three-dimensional models generated for AS1411, AS1411–8-17-769, and 8-17-769–AS1411, complexes formed between the Spike RNA fragment targeted by 8-17-769 and either the free deoxyribozyme or the two chimeric constructs were also modeled (Figure 7A–F). These models provided a structural framework for comparing the spatial organization of the aptamer and catalytic modules following conjugation and for assessing whether the interaction between the 8-17-769 region and the target RNA could be preserved within the chimeric context.

**Figure 7.**
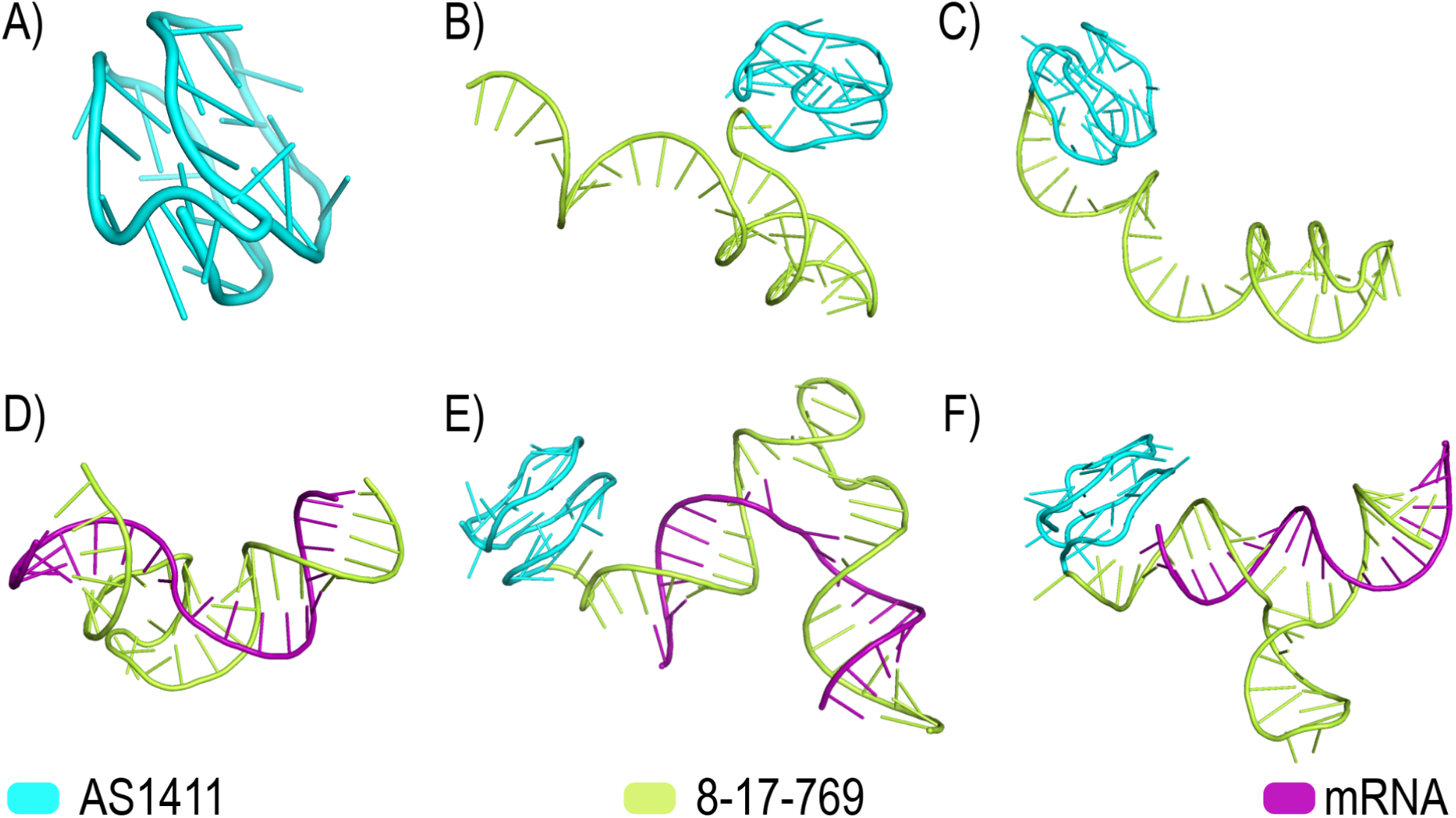
Three-dimensional oligonucleotide models generated using AlphaFold: **(A)** AS1411, **(B)** AS1411–8-17-769 conjugate, **(C)** 8-17-769–AS1411 conjugate, **(D)** 8-17-769–mRNA complex, **(E)** AS1411–8-17-769–mRNA complex, and **(F)** 8-17-769–AS1411–mRNA complex.

iMODS analyses of the RNA-containing complexes indicated that conjugation orientation may differentially affect the dynamic properties of the complexes (Figure 8A–C). In the free 8-17-769–RNA complex, localized regions of flexibility were observed along the catalytic deoxyribozyme, and this overall dynamic pattern was largely preserved in the 8-17-769–AS1411 conjugate (Figure 8A–B). In contrast, the AS1411–8-17-769 complex exhibited distinct deformability profiles, particularly in regions associated with the deoxyribozyme and the RNA. The preservation of the overall elastic network patterns suggests that both conjugates were capable of forming structurally compatible complexes with the RNA, whereas the orientation-dependent differences in deformability indicate that conjugation may alter local conformational flexibility. The three-dimensional models further showed that 8-17-769 and the target RNA could coexist within the same complex in both chimeric configurations (Figure 8C).

**Figure 8.**
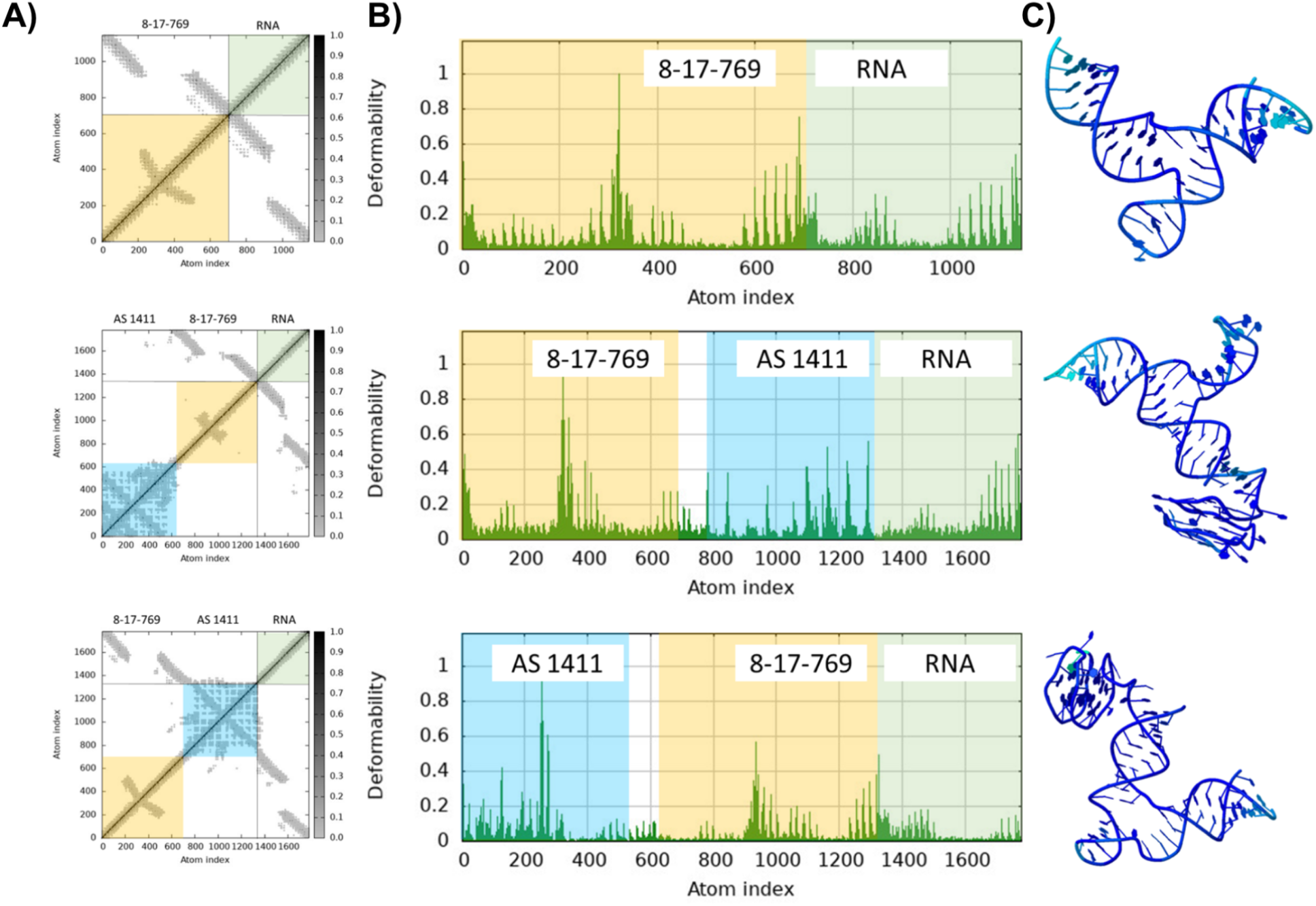
**C**omparison of the structural and dynamic properties of complexes formed between 8-17-769 and AS1411 conjugates and the target RNA. **A)** iMODS elastic network analyses of the 8-17-769–RNA, 8-17-769–AS1411–RNA, and AS1411–8-17-769–RNA complexes. The matrices represent patterns of collective motion and correlation among the atoms constituting the complexes. Colored regions indicate the corresponding oligonucleotide components. **B)** Deformability profiles of the same complexes calculated as a function of atom index. Higher peaks indicate regions with relatively greater flexibility or susceptibility to deformation within the complexes. **C)** Three-dimensional structural models of the analyzed complexes. From top to bottom, the panels show the 8-17-769–RNA, 8-17-769–AS1411–RNA, and AS1411–8-17-769–RNA complexes, respectively.

A second requirement for the chimeric design was that the AS1411 module retain a conformation compatible with interaction with its known molecular target, nucleolin, after conjugation to the deoxyribozyme. Therefore, the putative complexes of free AS1411 and both chimeras with nucleolin were modeled using HADDOCK 2.4. The HADDOCK score for the free AS1411–nucleolin complex was calculated as −146.5 ± 2.1, whereas the AS1411–8-17-769–nucleolin and 8-17-769–AS1411–nucleolin complexes yielded scores of −128.6 ± 10.1 and −152.1 ± 8.6, respectively (Table 3; Figure 10). The corresponding Z-scores were −2.1, −1.8, and −1.5, respectively.

**Figure 10.**
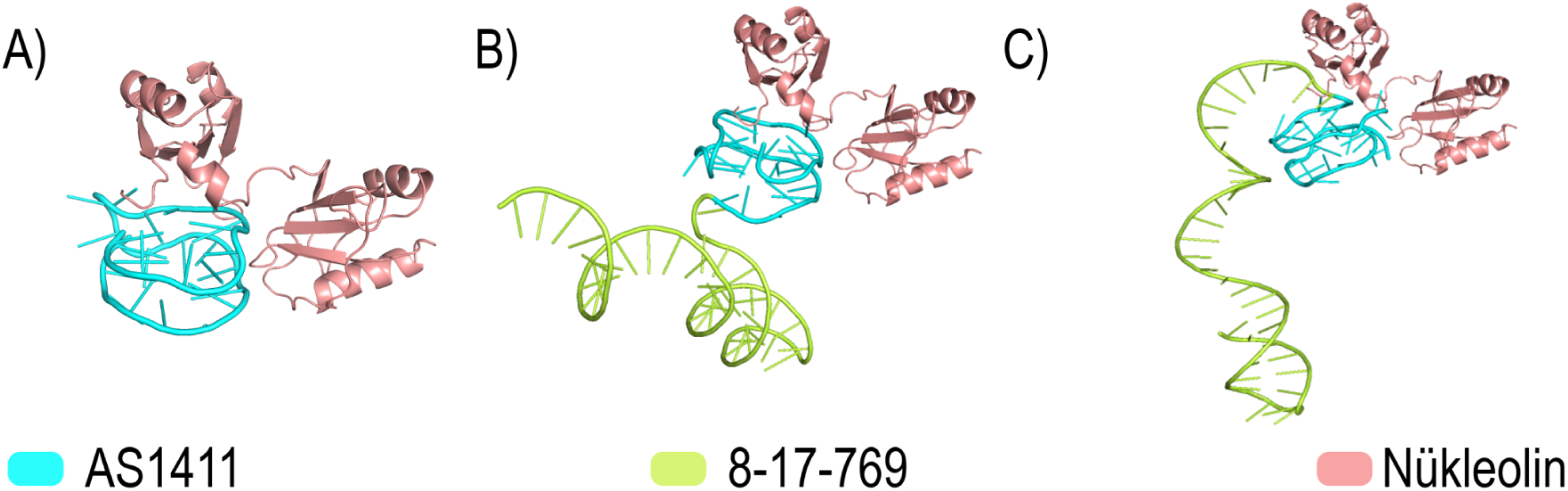
Three-dimensional models generated by protein–DNA molecular docking performed on the HADDOCK 2.4 web server: **(A)** AS1411–nucleolin complex, **(B)** AS1411–8-17-769–nucleolin complex, and **(C)** 8-17-769–nucleolin complex.

**Table 3.** Protein–DNA complexes modeled using the HADDOCK 2.4 web server, including corresponding HADDOCK scores and Z-scores.

| Complex | HADDOCK Score | Z-Score |
| --- | --- | --- |
| AS1411:Nucleolin | -146.5 (+/-) 2.1 | -2.1 |
| AS1411-8-17-769:Nucleolin | -128.6 (+/-) 10.1 | -1.8 |
| 8-17-769-AS1411:Nucleolin | -152.1 (+/-) 8.6 | -1.5 |

The ability to obtain docked nucleolin-containing complexes for both chimeras indicates that conjugation of AS1411 to the deoxyribozyme did not necessarily abolish the modeled interaction with nucleolin. Nevertheless, a clear difference was again observed between the two orientations: the 8-17-769–AS1411–nucleolin complex exhibited a more negative HADDOCK score than the AS1411–8-17-769 complex and was at least comparable to the value calculated for free AS1411 (Table 3). This observation provides additional in silico support for the possibility that, in the 8-17-769–AS1411 orientation, the AS1411 module retains an interaction geometry compatible with nucleolin binding.

However, it is important not to interpret HADDOCK scores as experimental binding affinities. Although these scores can be used to assess the relative structural and energetic favorability of the modeled complexes, they do not provide a direct *k_d_* value or a measure of cellular binding or internalization. Similarly, the Z-score should not be regarded as an independent measure confirming biological binding; rather, it reflects the relative position of a given HADDOCK cluster with respect to the other generated solutions. Therefore, the present molecular docking results should be interpreted not as direct evidence of AS1411-mediated cellular entry, but as supportive computational evidence indicating that structural configurations compatible with nucleolin interaction can be retained following conjugation.

When CD spectroscopy, computational analysis of the RNA-containing complexes, and nucleolin docking results are considered together, a consistent orientation-dependent pattern emerges. The 8-17-769–AS1411 chimera, which exhibited the stronger silencing profile in the cellular experiments, also showed a CD profile that better preserved the spectral characteristics of the individual components, displayed structural behavior in the RNA complex that more closely resembled the free deoxyribozyme system, and formed a docking model compatible with nucleolin interaction. In contrast, the AS1411–8-17-769 orientation exhibited more pronounced structural perturbation and reduced deformability in specific regions. Taken together, these parallel findings suggest that module order in aptamer–deoxyribozyme chimeras is not merely a design detail, but may represent an important design parameter capable of influencing the structural integrity of both the catalytic and targeting modules and, consequently, the overall functional performance.

Therefore, the overall prominence of 8-17-769–AS1411 in this study was not based on a single experimental measurement. Following the selection of 8-17-769 as the fastest RNA-cleaving candidate in the *in vitro* screening, conjugation of this catalytic module with AS1411 in the 8-17-769–AS1411 orientation was associated with the most pronounced reduction in S-gene expression in the cellular model. Structural and computational analyses further indicated that this configuration was consistent with an organization that induced less perturbation of the functional components. Taken together, these findings establish a mutually supportive experimental chain linking catalytic candidate selection, cellular function, and molecular structure.

## CONCLUSION

This study established a sequential experimental framework for the development of an aptamer–deoxyribozyme construct targeting SARS-CoV-2 Spike RNA, beginning with catalytic candidate selection and extending to cellular and structural evaluation. Comparative screening of eight deoxyribozymes revealed pronounced target-site-dependent differences in RNA cleavage and identified 8-17-769 as the lead candidate based on its rapid, reproducible cleavage profile and the highest observed cleavage rates among the tested sequences. Importantly, incorporation of this catalytic module into AS1411-containing chimeras demonstrated that functional performance was dependent not only on the presence of the two modules but also on their linear orientation. The 8-17-769–AS1411 configuration produced the greatest reduction in S gene expression in the cellular model and was simultaneously associated with a CD profile more closely resembling the individual components, preservation of deoxyribozyme-like dynamic behavior within the modeled RNA complex, and a docking configuration compatible with nucleolin interaction. These convergent findings suggest that conjugation orientation represents a critical design parameter in multifunctional oligonucleotide constructs, as module order may influence the structural integrity and accessibility of both catalytic and targeting domains. Although direct AS1411-mediated cellular uptake, experimental nucleolin-binding affinity, and antiviral activity in an infection model remain to be established, the present results provide a coherent basis for further development of 8-17-769–AS1411 and highlight the broader importance of structure-guided orientation optimization in aptamer–deoxyribozyme therapeutics.

## Supporting information

S-gene sequence

