## Supplementary material for "Identification of a SARS-CoV-2 Spike RNA-Cleaving DNAzyme and Optimization of Its AS1411 Chimera": S-gene sequence

S gene with T7 promoter

(5'-**TAATACGACTCACTATAGAAGCTTGG**ATGGAAAGTGAGTTCAGAGTTTATTCT
AGTGCGAATAATTGCACTTTTGAATATGTCTCTCAGCCTTTTCTTATGGACCTTGAA
GGAAAACAGGGTAATTTCAAAAATCTTAGGGAATTTGTGTTTAAGAATATTGATGG
TTATTTTAAAATATATTCTAAGCACACGCCTATTAATTTAGTGCGTGATCTCCCTCAG
GGTTTTTCGGCTTTAGAACCATTGGTAGATTTGCCAATAGGTATTAACATCACTAGG
TTTCAAACTTTACTTGCTTTACATAGAAGTTATTTGACTCCTGGTGATTCTTCTTCA
GGTTGGACAGCTGGTGCTGCAGCTTATTATGTGGGTTATCTTCAACCTAGGACTTT
TCTATTAAAATATAATGAAAATGGAACCATTACAGATGCTGTAGACTGTGCACTTGA
CCCTCTCTCAGAAACAAAGTGTACGTTGAAATCCTTCACTGTAGAAAAAGGAATC
TATCAAACTTCTAACTTTAGAGTCCAACCAACAGAATCTATTGTTAGATTTCCTAAT
ATTACAAACTTGTGCCCTTTTGGTGAAGTTTTTAACGCCACCAGATTTGCATCTGTT
TATGCTTGGAACAGGAAGAGAATCAGCAACTGTGTTGCTGATTATTCTGTCCTATAT
AATTCCGCATCATTTTCCACTTTTGGATCCCGG-3')

*Bold font, T7 promoter region.
